# A New Fight-or-Flight Pacemaker Mechanism via Ryanodine Receptor Abundance and Superclustering

**DOI:** 10.1101/2025.11.26.690788

**Authors:** Valeria Ventura Subirachs, Syevda Tagirova, Alexander V Maltsev, Dongmei Yang, Edward G Lakatta, Michael D Stern, Victor A Maltsev

## Abstract

The sinoatrial node is the primary cardiac pacemaker. Individual sinoatrial node cells (SANCs) generate spontaneous rhythmic action potentials (APs) that initiate each heartbeat. The mechanism of SANC automaticity and its modulation by autonomic nervous system are based on the coupled function of molecules of both the cell membrane (ion channels, exchangers, and pumps) and the sarcoplasmic reticulum (SR), which generates rhythmic local Ca releases (LCRs). While LCRs are generated by ryanodine receptors (RyRs), the molecular-scale RyR network structure remains unknown. Here we performed single-molecule localization of RyRs via direct Stochastic Optical Reconstruction Microscopy (dSTORM) in rabbit SANCs in basal conditions and 5 minutes after β-adrenergic receptor (βAR) stimulation by isoproterenol. RyRs form clusters of various sizes, with a mean density of 67.7±13.2 RyR/μm^2^. (Mean±SEM, 6 cells). While the majority of cluster sizes ranged from 3 to 32 RyRs, each cell had a few substantially larger clusters (>76 RyRs), dubbed superclusters. βAR stimulation significantly increased the RyR density to 119.1±22.6 RyR/μm^2^ (8 cells, p<0.05) and created more superclusters. Our new numerical SANC model showed that superclustering substantially decreased the AP cycle length (APCL) by creating Ca release hotspots that initiated larger LCRs under any condition. Increasing RyR density prolonged APCL in the basal state but shortened APCL during βAR stimulation. With no change in RyR network, βAR stimulation of only SR Ca pump and ion currents shortened APCL on average from 414.9 to 284 ms. When realistic higher RyR density and superclustering were added to the model, APCL further shortened to 231.9 ms. Thus, dynamic nanoscale changes in RyR network provide a new powerful pacemaker mechanism. This mechanism may help explain athletic bradycardia at rest and high exercise heart rates, with its deterioration contributing to the age-related loss of heart rate reserve.

**Author summary:** The heartbeat begins in the sinoatrial node, the heart’s natural pacemaker. During stress or exercise, pacemaker cells accelerate their firing rate via β-adrenergic (“fight-or-flight”) stimulation. Although Ca signals are known to drive this acceleration via electrogenic Na/Ca exchanger, how the underlying ryanodine receptor (RyR) Ca release channels are organized at the nanoscale and how this organization changes during stimulation remain unclear. Here, we used super-resolution microscopy (dSTORM) to visualize individual RyRs in rabbit pacemaker cells. We found that RyRs form heterogeneous clusters that include rare, very large “superclusters.” β-adrenergic stimulation increased overall RyR density, enlarged clusters, and markedly increased the number of superclusters. To understand how these structural changes affect pacemaker function, we developed a computational sinoatrial node cell model in which each RyR was represented explicitly and arranged according to our imaging data. Numerical model simulations revealed that superclusters act as Ca-release hotspots that fire early during diastolic depolarization and recruit neighboring RyR clusters to fire via Ca-induce-Ca-release mechanism. When combined with β-adrenergic enhancement of Ca cycling and membrane currents, the superclustering and RyR abundance synergistically accelerated action potential firing. Together, our results identified dynamic nanoscale RyR network reorganization as a key structural mechanism in fight-or-flight response.

## Introduction

The sinoatrial node (SAN) serves as the heart’s primary pacemaker, with individual sinoatrial cells (SANCs) generating spontaneous, rhythmic electrical signals, action potentials (APs) that initiate each heartbeat [1, 2]. The underlying cellular mechanism driving this automaticity includes a complex interplay of multiple signals within a coupled-clock system [3, 4], involving molecules on the cell surface membrane (ion channels, exchangers, and pumps), known as a membrane clock, and internal Ca-handling structures, primarily the sarcoplasmic reticulum (SR), known as a Ca clock. The Ca clock has two major components: (i) a Ca pump (SERCA molecules) that pumps Ca from cytoplasm into the SR and (ii) Ca release channels (ryanodine receptors, RyRs) that spontaneously rhythmically release Ca from the SR via local Ca releases (LCRs). The LCRs are complex locally propagating events of different morphologies [5] generated by RyR clusters (also known as Ca release units, CRUs) forming an intricate intracellular network interacting via Ca-induced-Ca-release (CICR) mechanism.

The LCRs contribute significantly to the pacemaker rate via activation of electrogenic Na/Ca exchanger (NCX) that timely depolarizes the cell membrane [6, 7]. This LCR-NCX-induced membrane depolarization activates near-threshold L-type Ca channels, especially the low-voltage-activated Cav1.3 isoform [8], that generate more depolarization and Ca influx that activates more LCRs, forming a positive feed-back loop, known as AP ignition [9]. Thus, the RyR network, especially near the cell surface membrane, where RyRs interact with membrane proteins, is crucially important for the generation, synchronization, and propagation of LCRs that regulate the SANC pacemaker function.

The distribution of RyRs in SANCs has been extensively studied by different imaging techniques and numerical modeling. Our previous studies showed that i) synchronization of stochastic CRUs creates a rhythmic Ca clock, and the transition in LCR characteristics is steeply nonlinear over a narrow range of release current, resembling a phase transition [10]; ii) RyR-NCX-SERCA local crosstalk ensures pacemaker cell function at rest and during the fight-or-flight reflex [11], iii) hierarchical clustering of RyRs is important for CICR propagation [12]; iv) CICR facilitation by heterogeneities in CRU sizes and locations regulates and optimizes cardiac pacemaker cell operation under various physiological conditions [13, 14]. While previous studies showed functional importance of fine structure of the RyR network, the RyR cluster locations and sizes were imaged by confocal microscopy [12, 15, 16] and more recently by structured illumination microscopy [14] which cannot resolve individual RyR channels. On the other hand, previous electron microscopy studies did resolve individual RyRs located in peripheral couplings in SANCs [17], but did not describe the RyR network in sufficient detail to glean further functional insight.

Direct Stochastic Optical Reconstruction Microscopy (dSTORM) is a new super-resolution imaging technique capable of localizing individual molecules with approximately 20 nm resolution [18]. Interplay between RyR arrangement is important for control of Ca release in health and disease (review [19]). Individual RyRs have been previously detected by dSTORM only in cardiac ventricular cells [20, 21] showing that the RyR network is dynamic [22–25] and that RyR clusters expand and coalesce after application of isoproterenol, a β-adrenergic receptor (βAR) agonist simulating fight-or-flight response [24].

To further understand how RyRs contributes to cardiac pacemaker function, two major topics require clarification: 1) RyR network structure at the molecular scale in SANC and 2) the network changes during βAR stimulation and their contribution to AP firing regulation. To address this knowledge gap, we used dSTORM-based single molecular localization to examine the RyR distribution in rabbit SANCs, in cell periphery close to the cell membrane. We found that RyR channels are not uniformly distributed but have a few substantially larger clusters, dubbed superclusters. βAR stimulation significantly increased the RyR density and created more superclusters. To obtain functional insights, we performed numerical model simulations using a new computational SANC model featuring single RyR resolution and incorporating our new experimental results. Our simulations showed that RyR abundance and superclustering is a new powerful pacemaker mechanism; and that the strongest βAR stimulation effect is achieved when all pacemaker mechanisms work together, i.e. increases in RyR density and superclustering are matched with respective increases in SR Ca pumping and membrane ion currents (I_CaL_, I_f_, and I_Kr_).

## Results

### βAR stimulation increases RyR density and cluster size and shorten nearest neighbor distances

We measured and analyzed 14 SANCs, comprising 6 cells in the basal state and 8 cells under βAR stimulation. RyRs form clusters of various sizes, with the mean density of RyRs of 67.7±13.2 RyR/μm² (Mean±SEM) in SANCs in the basal state. βAR stimulation significantly increased the RyR density to 119.1±22.6 RyR/μm² (p<0.05) and created more superclusters. As summarized in S1 Table, βAR stimulation induced significant structural changes in the RyR network. RyR density increased by 76.1% (to 119.11 RyR/μm^2^). Mean cluster size increased by 30.2% (from 17.40 to 22.66 RyR/cluster). Concurrently, the network became more compact, with the mean nearest neighbor distance between clusters decreasing by 34.5% (from 180.15 nm to 118.02 nm). These comparisons are visualized in the box plots in Fig 1. All metrics showed statistically significant differences between the basal and βAR groups (two-sided Mann-Whitney U test, p < 0.05)

**Fig 1.**
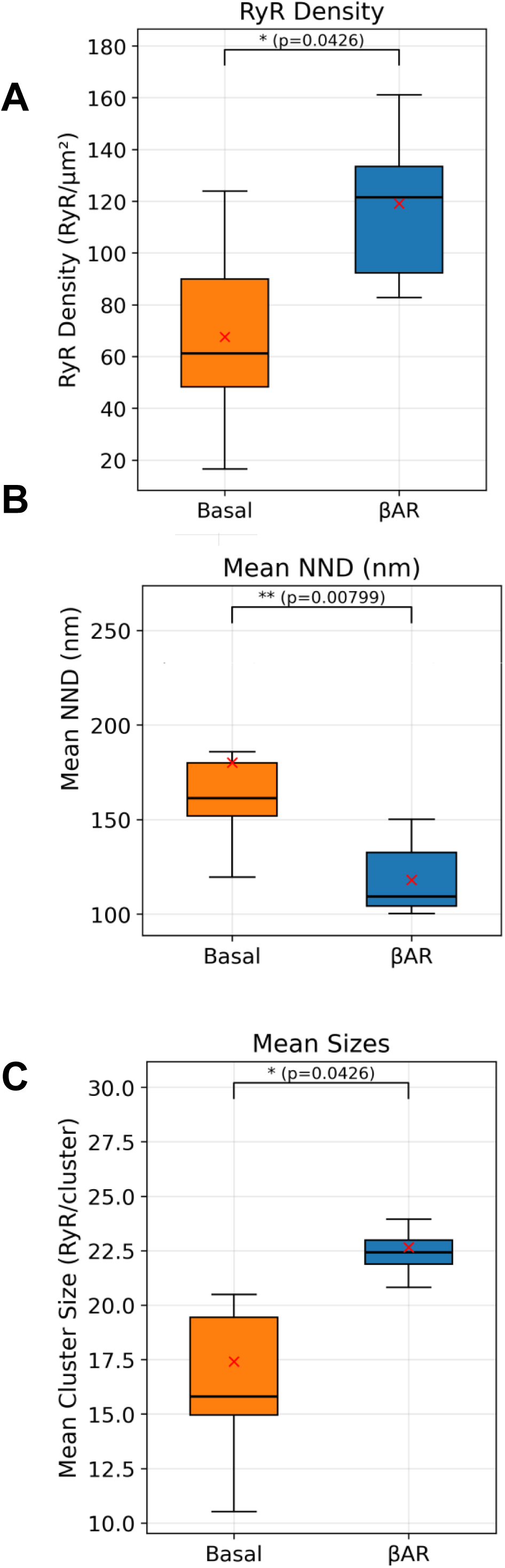
Box-plot comparisons of RyR density, spacing, and cluster size between basal and βAR stimulation. Presented are per-cell metrics from S1 Table derived from DBSCAN-labeled localizations. **A–D.** Box plots show RyR density RyR/μm^2^), mean nearest neighbor distance (NND, nm) and mean cluster size (RyR/cluster). Box plots display the median (solid horizontal line), interquartile range (box), whiskers extending to 1.5× the interquartile range. Cross shows mean values. P-values above each panel are from two-sided Mann–Whitney U tests; all comparisons shown are significant (p ≤ 0.05). Together, panels A–D indicate significantly increased density and larger clusters with shorter cluster spacing under βAR stimulation. Cells were treated with 0.3 µM isoproterenol for 5 min at 37 °C; control cells received vehicle only.

### βAR stimulation increases RyR supercluster number and sizes

We further examined the increase in cluster size by analyzing the cluster size distributions (Fig 2). In a representative cell in the basal state (Cell 6, Fig 2A), the distribution is highly skewed, with a rapid decay from the smallest cluster sizes. In contrast, the βAR-stimulated cell (Cell 12, Fig 2C) exhibits a right-shifted and heavier tail, indicating a significantly higher population of large aggregates. Panels B and D in Fig 2 show examples of RyR distributions in cell regions in basal state and in the presence of βAR stimulation that clearly visualize the change. While under the basal state, cells contained large clusters (e.g., 99 and 185 RyR), the βAR-stimulated cells featured “superclusters” that were not prevalent in the basal state, reaching sizes up to 414 RyR. This indicates that βAR stimulation promotes the formation of markedly larger RyR aggregates. During βAR stimulation, the supercluster size significantly (p<0.05) increased by 52.4%, from 54.09 to 82.43 RyR/cluster (see inset in Fig 2C).

**Fig 2.**
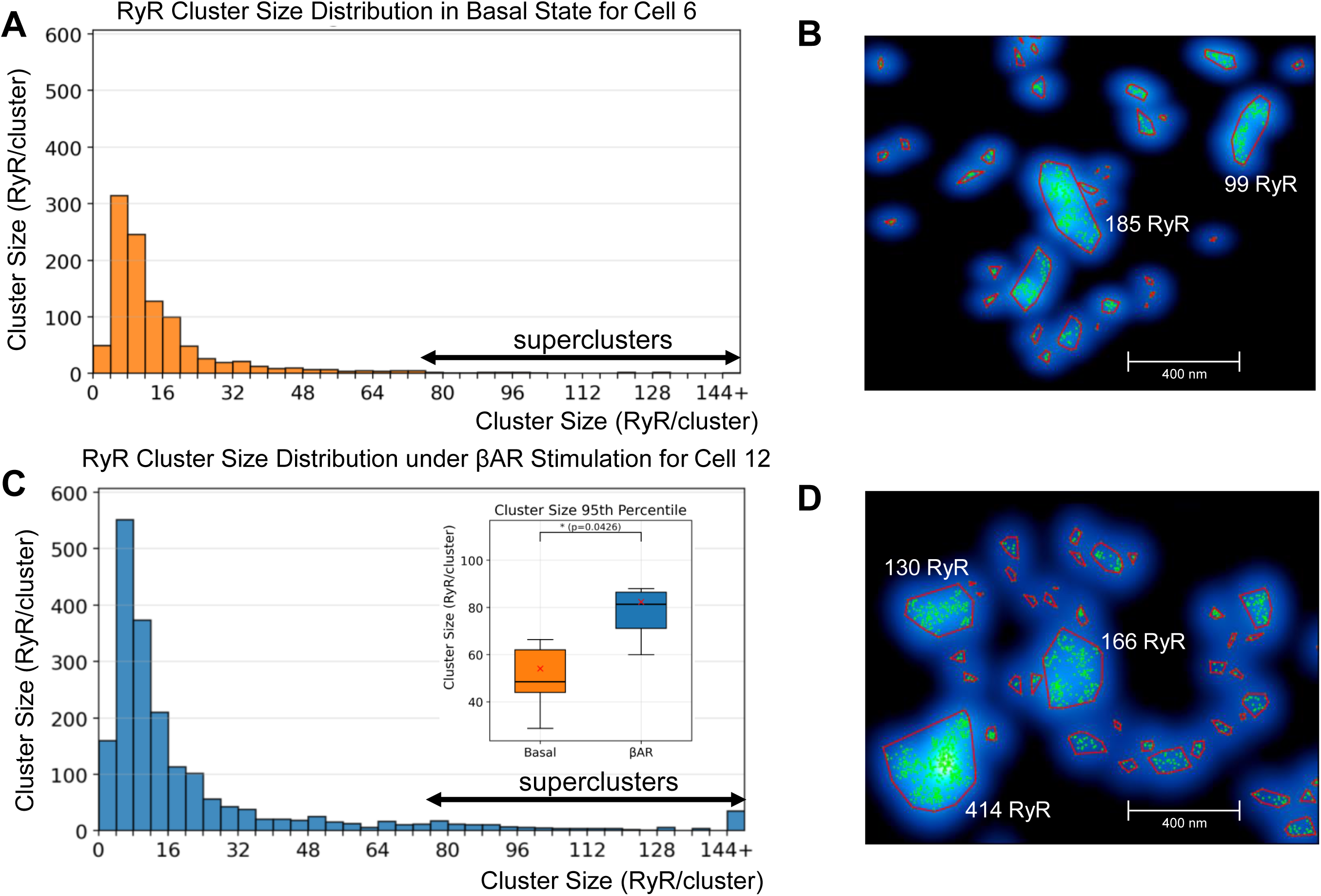
RyR cluster-size distributions and exemplary large clusters underβAR stimulation. Presented are DBSCAN-derived cluster sizes (RyR per cluster) and representative regions from basal and βAR cells. **A, C.** Histograms of cluster size for a basal cell (Cell 6) and a βAR-stimulated cell (Cell 12) show more clusters overall and a right-shifted, heavier tail under βAR, indicating larger aggregates. Double headed arrows show superclusters with sizes >76 RyRs as defined in the text. **B, D.** Examples of the largest clusters in each condition overlaid on a smoothed density map (blue glow); red polygons mark cluster boundaries, numbers indicate RyR per cluster. The basal field contains large clusters (e.g., 185 and 99 RyR), whereas βAR exhibits superclusters reaching >400 RyR. The numbers of clusters and cluster sizes increased during βAR stimulation. Inset in Panel C shows effect of βAR stimulation on cluster-size within 95th percentile (RyR/cluster) in a box plot. P-value (0.0426) is from two-sided Mann–Whitney U test (see numerical values in S1 Table).

### βAR stimulation changes spatial organization within RyR clusters

RyR clusters were categorized into size groups based on the pooled distribution of cluster sizes across all cells and conditions. We first identified all clusters from 14 cells (6 basal, 8 βAR-stimulated) at the 5-minute timepoint and ranked them by the number of RyRs per cluster. Size categories were defined using percentile cutoffs from this combined distribution:

- Small: ≤6 RyRs (≤25th percentile)
- Medium-Small: 7-10 RyRs (25th-50th percentile)
- Medium-Large: 11-21 RyRs (50th-75th percentile)
- Large: 22-76 RyRs (75th-95th percentile)
- Supercluster: >76 RyRs (>95th percentile), see examples shown by double headed arrows in Fig 2A and C.

This percentile-based approach provides biophysically meaningful categories (as we show below in our SANC model simulations) that capture the natural distribution of cluster sizes while maintaining sufficient sample sizes for statistical analysis.

To quantify the internal spatial organization of RyRs within individual clusters, we applied the Hopkins statistical analysis (results are summarized in Fig 3 and mathematical details in Methods section). This analysis revealed size-dependent changes in internal RyR organization following β-adrenergic stimulation. Medium-Large clusters (11-21 RyRs) showed a modest but highly significant increase in Hopkins values (1.8% change, p<0.001), though values remained near 0.5 in both conditions, indicating maintenance of near-random internal organization. In contrast, Superclusters (>76 RyRs) demonstrated a pronounced increase in Hopkins values (6.2% change, p<0.01), though absolute values remained below 0.4, suggesting these large clusters maintain relatively uniform RyR distribution despite βAR-induced reorganization.

**Fig 3.**
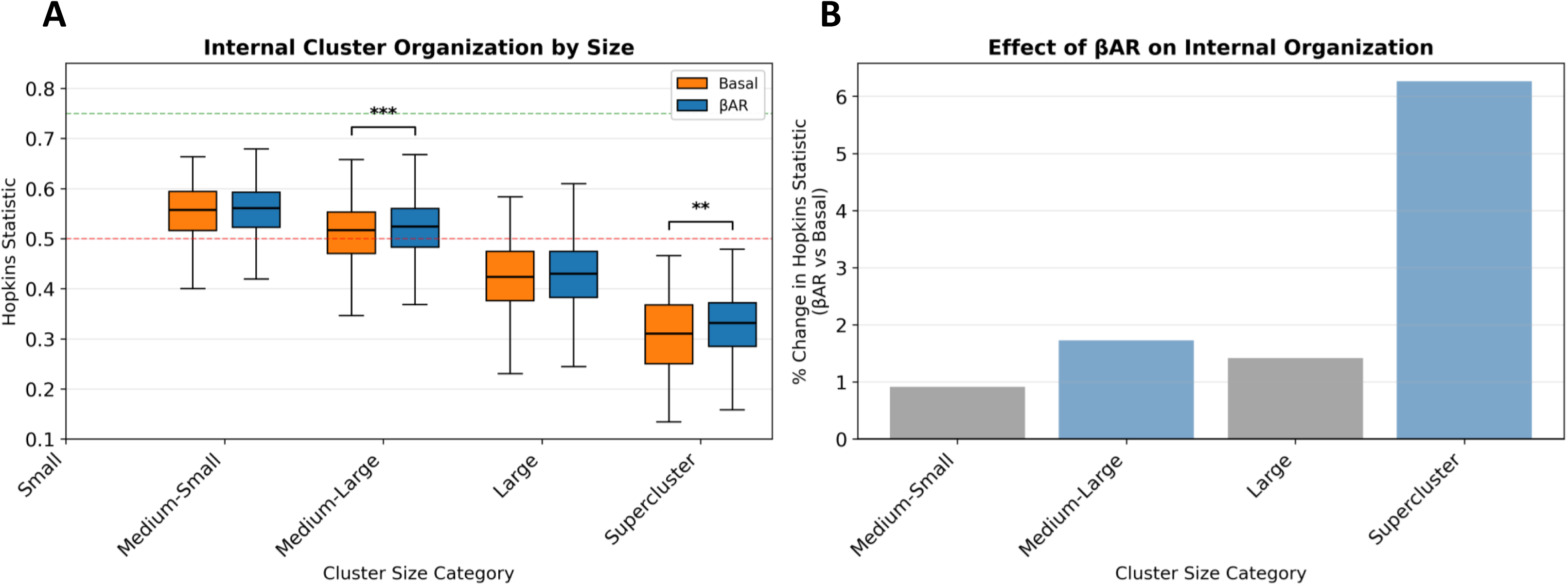
Hopkins statistic analysis reveals size-dependent changes in internal RyR organization followingβ-adrenergic stimulation. **A.** Box plots showing Hopkins statistic values for RyR clusters grouped by size category under basal (orange) and βAR-stimulated (blue) conditions. Horizontal brackets with asterisks indicate statistically significant differences between Basal and βAR Hopkins statistic distributions groups in the Medium-large and Supercluster categories (Mann-Whitney U test; **p<0.01, ***p<0.001). Hopkins values near 0.5 (red dashed line) indicate random distribution, values <0.5 indicate regular/uniform spacing, and values approaching 0.75 (green dashed line) indicate highly clustered internal organization. **B.** Percent change in Hopkins statistic following βAR stimulation for each cluster size category. Data from n=6 basal and n=8 βAR-stimulated cells containing 275, 1440, 939, and 174 clusters (basal) and 474, 2729, 2369, and 689 clusters (βAR) for Medium-Small, Medium-Large, Large, and Supercluster categories, respectively.

The lower Hopkins values observed in superclusters (0.31-0.33) compared to smaller cluster categories indicate more uniform spacing of RyRs within these large functional units. The βAR-induced increase in Hopkins values for superclusters, while maintaining sub-0.5 values, suggests a transition from highly regular to moderately regular organization, potentially creating functional sub-domains within the cluster. This size-dependent reorganization aligns with the functional requirement for different cluster sizes to serve distinct roles in cardiac pacemaking, with larger clusters requiring more sophisticated internal architecture to coordinate Ca release across numerous RyRs.

### Structure-function relationship of RyR network revealed by numerical model simulations

To further interpret our experimental findings and obtain functional insights into how RyR network regulate pacemaker function, we performed numerical model simulations. We simulated SANC function for eight scenarios representing all possible combinations of 2 densities (normal and high), 2 RyR clustering (normal clustering and superclustering) and 2 coupled-clock parameter sets, i.e. P_up_ (SR Ca pump), I_CaL_, I_Kr_ and I_f_ for basal state and βAR stimulation. Normal (low) RyR density was set to 67.65 RyR/μm^2^, the average density in the basal state and high density was set to 119.07 RyR/μm^2^, the average density during βAR stimulation (S1 Table). Normal clustering was set as measured in Cell 6 (basal state) and superclustering was set as measured in Cell 12 (βAR stimulation). Results of the simulations are presented in Fig 4. Representative examples of submembrane Ca dynamics together with functional state of each CRU (color coded) are shown in S1-S8 Videos for each scenario.

**Fig 4.**
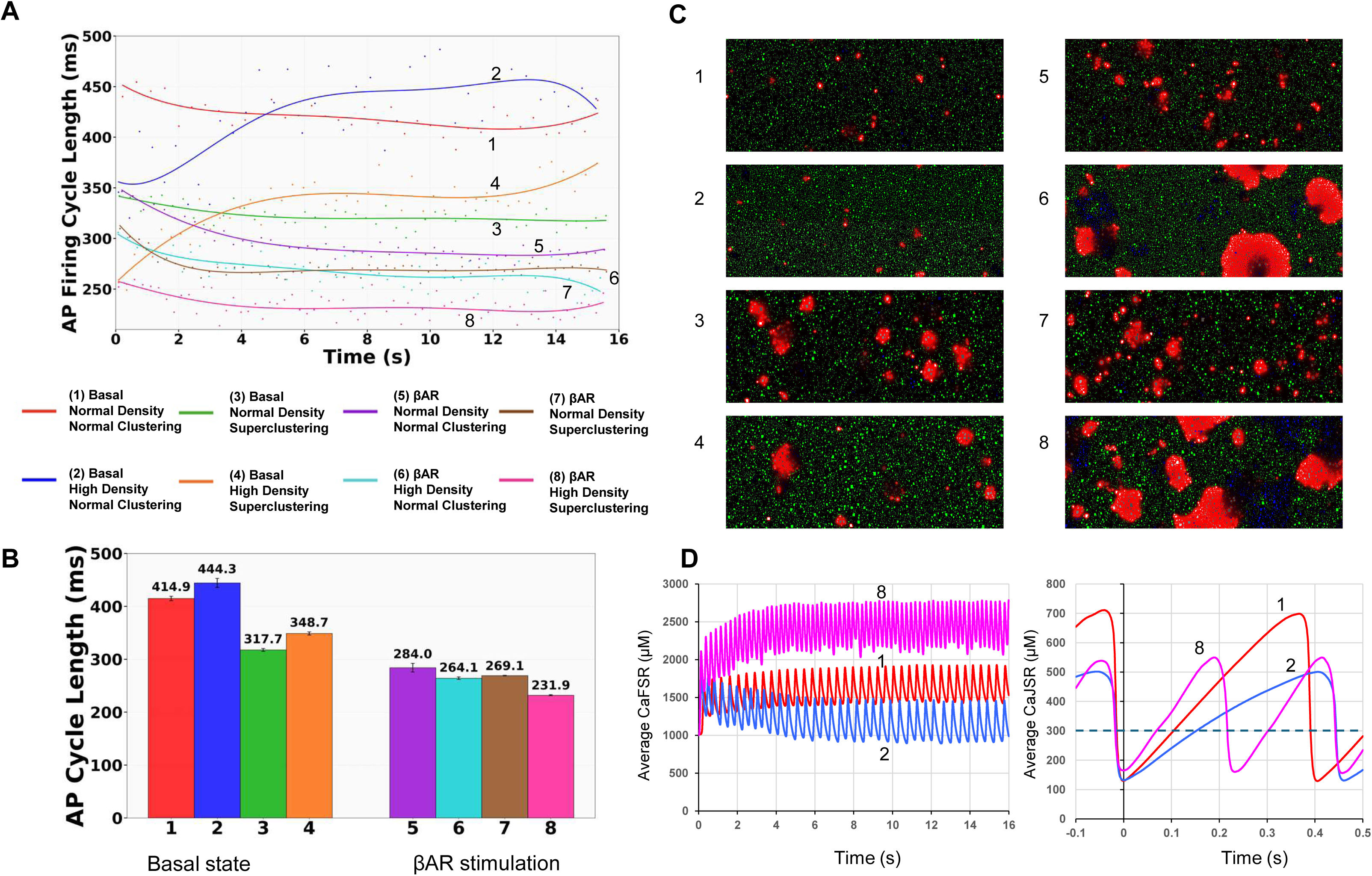
Synergistic effect of RyR density and superclustering on AP firing cycle length duringβAR stimulation revealed by numerical model simulations. Shown are simulations for 8 different scenarios (shown by respective numbers at the individual plots and labels) performed for combinations of i) two different RyR densities found at the basal state (in scenarios labeled “normal density” of 67.65 RyR/μm^2^) and during βAR stimulation (labeled “high density” of 119.07 RyR/μm^2^); ii) two cluster-size distribution found in the basal state (labelled “Normal clustering” taken form Cell 6, Fig 2A) and during βAR stimulation (labelled “superclustering” taken from Cell 12; Fig 2C); and iii) for basal state or βAR stimulation (labelled “Basal” or “βAR”, respectively). **A.** APCL versus time in all 8 scenarios; solid curves show respective polynomial fits. **B.** Bar plots summarize the steady-state APCL (mean±SD) measured wthin 6 s to 16 s in five independent runs for each scenario. (see also S1-S8 Videos). (**C**) Snapshots of S1-S8 Videos showing submembrane local Ca dynamics and CRUs at −45 mV near I_CaL_ activation threshold for each simulation scenario labeled by numbers at the snapshots. [Ca] was coded by red shades from black (0.15 μM) to pure red (>10 μM). Refractory CRUs are shown in blue shades; CRUs ready to release are in green; and CRUs releasing Ca are in grey shades. Both blue shades and white shades reflect JSR Ca changes, with a saturation level set at 0.3 mM. **D.** Dynamics of Ca concentration in free SR (CaFSR, left panel) and junctional SR (CaJSR, right panel) in scenarios 1, 2 and 8. In the right panel, the time was shifted to illustrate and compare different rates of junctional SR refilling with Ca from the nadir (t=0). Horizontal dashed line shows the Ca spark activation threshold (300 μM) in our model.

Simulations revealed that increasing RyR density alone surprisingly prolonged the AP cycle length (APCL) in the basal state firing but shortened APCL during βAR stimulation. However, introducing superclustering alone shortened APCL under both basal and during βAR stimulation. The combination of high density and superclustering, representing the full βAR phenotype that we found experimentally, resulted in a synergistic effect that yielded the shortest APCL of all conditions (231.9 ms, magenta color in Fig 4). The critical moment in clock coupling happens close to the threshold of I_CaL_, near the AP ignition onset when LCR activity increases [9]. To get insights into the coupling process, for each scenario we provided here snapshots (Figure 4C) of S1-S8 Videos showing submembrane Ca dynamics (coded in red shades) at −45 mV, reflecting LCR activity and their self-organization at that critical moment (Fig 4C). The LCR abundance in the snapshots was reversely linked to APCL: the lowest LCR activity was linked to longest APCL (slowest AP firing) in basal state with high RyR density and normal clustering (scenario 2), whereas the highest LCR activity was linked to the shortest APCL (fastest AP firing) in during βAR stimulation with high RyR density and superclustering (scenario 8). The LCR activity in scenario 8 was self-organized into large and abundant propagating events (S8 Video).

Further insights into different LCR activity in different RyR networks were obtained by examining dynamics of Ca concentration in free SR (CaFSR) and junctional SR (CaJSR). Both systolic and diastolic CaFSR levels substantially decreased in scenario 2 with increased RyR density (vs. scenario 1) but substantially increased in scenario 8 with the shortest APCL (Fig 4D, left panel). The CaFSR level decreased in denser RyR network in the basal state, because overall larger junctional SR (accommodating larger RyR numbers) with more connections to free SR shifts source-to-sink balance towards sink. The lower Ca loading of the free SR, in turn, resulted in slower refilling junctional SR and therefore delayed LCR occurrence during diastolic depolarization as reflected in rare small LCRs in scenario 2 (Fig 4C, right panel).

To learn more about functional importance of different size clusters, we collected statistics on cluster firing in each size category with respect to membrane potential at which they fire (Fig 5). Histograms of firing events at each potential showed that in the basal state (left panels) a large fraction of smaller clusters began firing at the rapid AP upstroke, i.e. at Vm>-20 mV. Thus, this fraction of smaller clusters stays in reserve and does not contribute to diastolic depolarization. In contrast, a major fraction of larger clusters (Medium-large, Large, and Superclusters) fired during diastolic depolarization (from −65 to −40 mV). Superclusters, being infrequent (>95th percentile), rigorously recruited (via CICR) their neighbors within a 1 μm neighborhood to fire as shown by orange histograms in each size category which were significantly shifted towards lower potentials (almost coinciding with the distribution of superclusters). During βAR stimulation (right panels) firing all clusters substantially shifted towards diastolic depolarization, indicating that those clusters that were in reserve during basal state now contribute to pacemaker function, representing a new mechanism of βAR stimulation.

**Fig 5.**
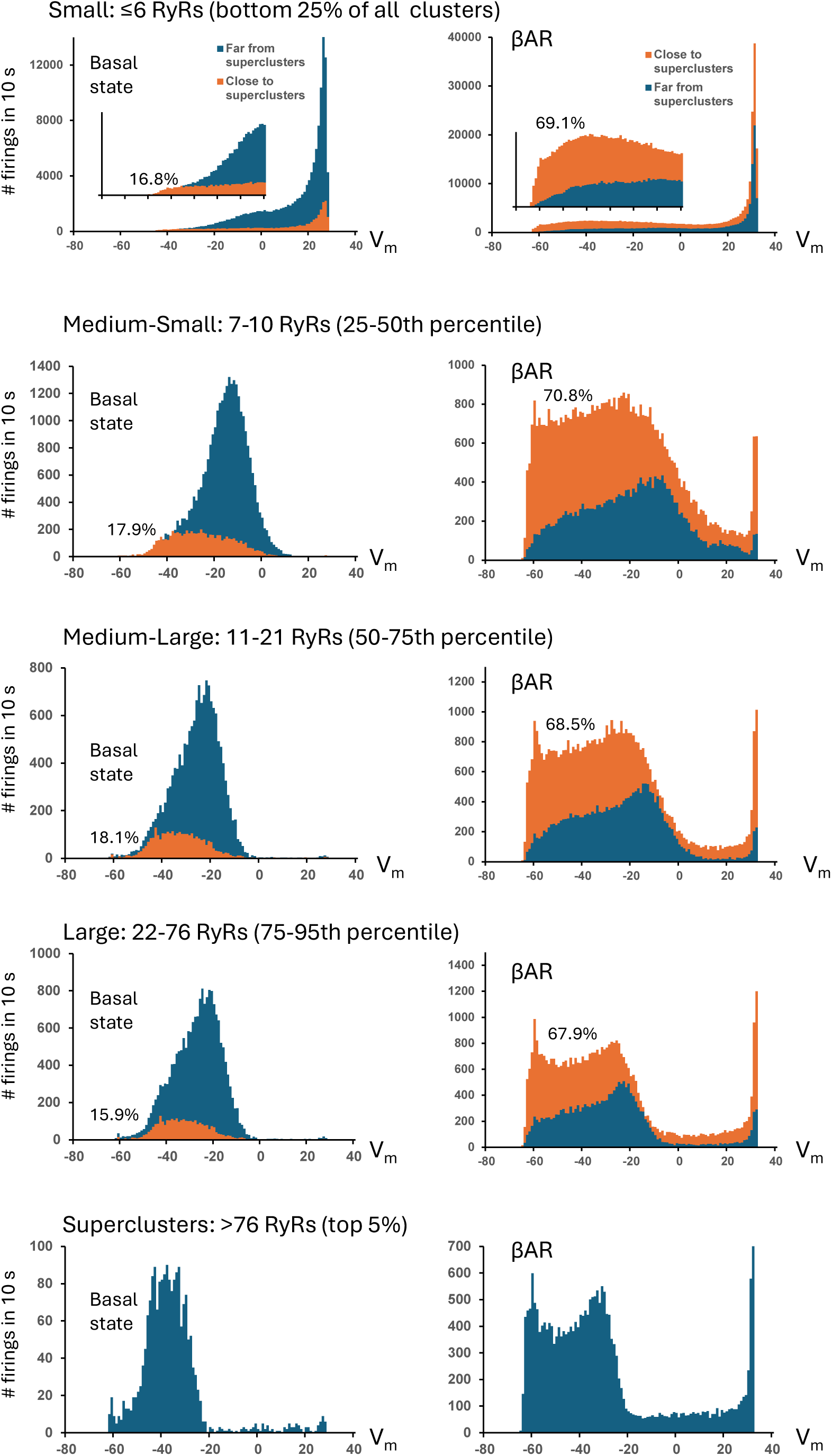
A new mechanism of βAR stimulation via CRU reserve and superclustering revealed by numerical model simulations of SANC function. Shown are results of statistical analysis of numerical simulations of SANC function in basal state (left panels) and during βAR stimulation (right panels) in each CRU size category with respect to membrane potential at which CRUs fire. The analysis was performed separately for CRUs located close superclusters (with 1 μm neighborhood, orange histograms) and far from the superclusters (dark blue histograms, overlapped). The orange histograms for CRUs near superclusters were shifted to lower membrane potentials (towards diastolic depolarization range) indicating their strong recruitment by superclusters. During βAR stimulation all histograms shifted towards diastolic depolarization range and percentage of CRUs close to superclusters (shown in each histogram) increased resulting in a stronger recruitment of CRUs by the superclusters especially at the very early stage of diastolic depolarization close to maximum diastolic potential near −65 mV.

How do superclusters participate in this mechanism? First, the network density increases during βAR stimulation, and therefore more clusters fall into the 1 µm neighborhood of the superclusters (67-71% during stimulation vs 16-18% in basal state, Fig 5). Next, Ca releases from clusters become stronger (with higher pumping rate, P_up_), especially in superclusters having more RyRs. Thus, overall recruitment capability of superclusters via CICR substantially increases. This results in a strong recruitment of neighboring RyRs to fire during early diastolic depolarization, creating a strong “jump” of orange histogram near maximum diastolic potential (MDP, near −65 mV). Other clusters (far from superclusters) fire stronger but at later stages of diastolic depolarization and during AP upstroke.

We further explored the isolated effect of RyR density while fixing the cluster-size distribution pattern as either in basal state or during βAR stimulation (Fig 6). Under basal conditions, increasing density notably prolonged APCL (from ∼375 ms to 429 ms). Conversely, during βAR-stimulation, increasing density shortened APCL (from ∼276 ms to 240 ms), accelerating the AP firing rate. This demonstrates that denser CRU networks work synergistically with upregulated Ca cycling, including SR pumping rate and increased membrane currents (βAR state) to shorten diastolic depolarization and increase the AP firing rate (Fig 6).

**Fig 6.**
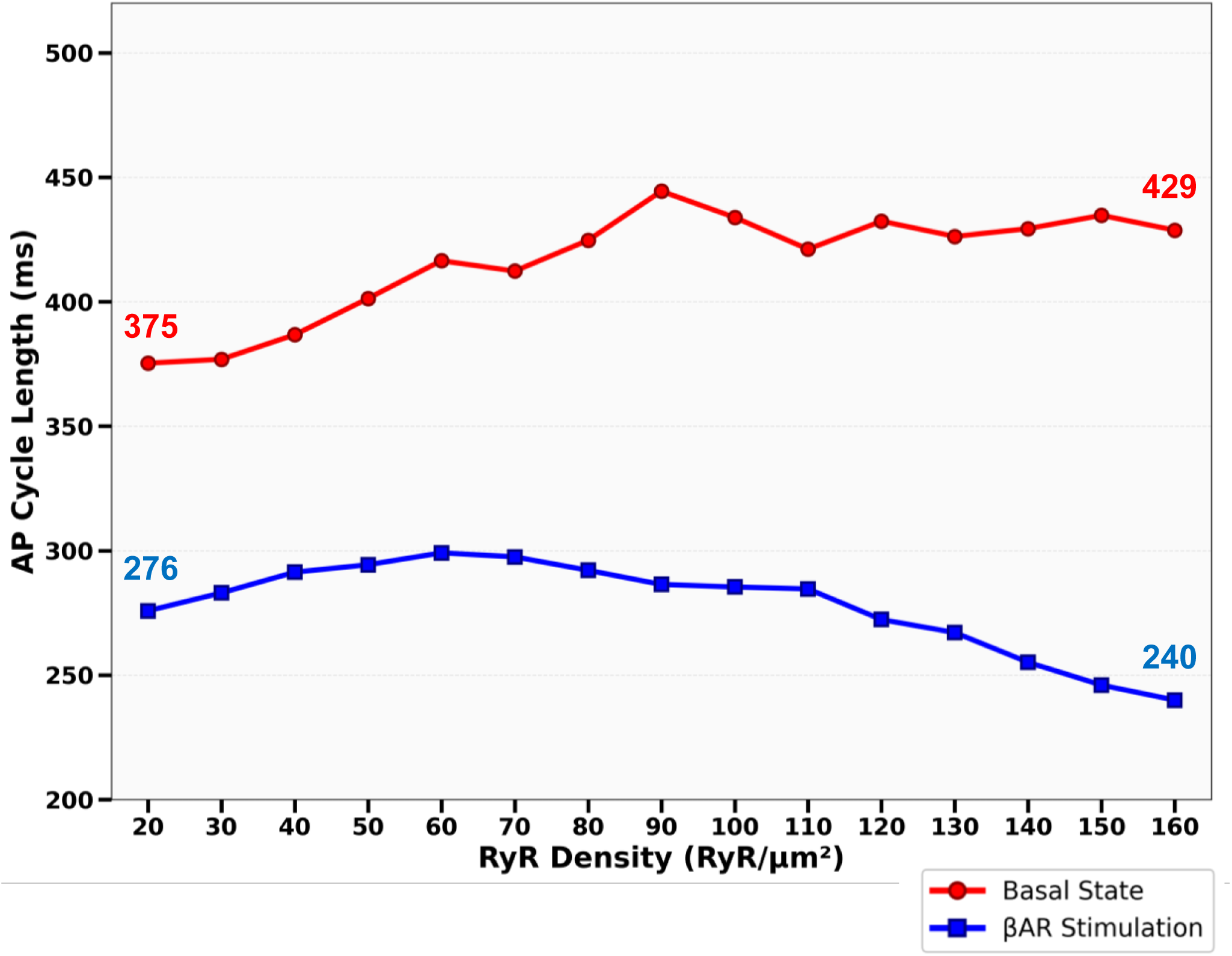
Dependence of AP cycle length on RyR surface density revealed by our numerical model simulations. Shown are steady-state AP cycle lengths (ms) simulated at basal (red) and βAR (blue) settings while varying RyR density from 10 to 160 RyR/μm^2^with a cluster-size distribution assigned for both βAR stimulation and basal state (Fig 2A).

## Discussion

Using dSTORM-based super resolution imaging combined with numerical modeling,we established the molecular-scale structure of the RyR network and discovered a novel fight-or-flight pacemaker mechanism via dynamic changes in RyR network. First, we quantified the network structure in basal and βAR-stimulated SANCs (Figs 1 and 2, S1 Table). We found that βAR stimulation substantially reorganizes the RyR network. RyR density markedly increases, while the nearest neighbor distances between clusters simultaneously decrease and mean cluster size increases. Importantly, the size increase was not uniform; it was characterized by the formation of “superclusters” (Fig 2), which produced a right-skewed size distribution with an extended upper tail. This reorganization was most evident in the 95th percentile of the cluster size distribution, which increased by 52.4%. Based on this data we developed a new numerical SANC model that can simulate the function of CRUs with single RyR resolution. Our model simulations showed that the strongest βAR stimulation effect is achieved when increases SR Ca pumping and membrane ion currents are combined with respective synergistic changes in RyR density and superclustering.

The functional expression of RyRs together with SR Ca pumping defines the capability of the Ca clock to generate stronger Ca releases. So far, the best estimate of about 70 channels per μm^2^ of cell surface has been based on a numerical comparison of immunofluorescence intensity of RyR distribution in serial sections of SANCs and ventricular cells [12]. This was an indirect estimate, whereas the direct measurements of the channel density have not been performed. We not only measured RyR density, but also performed numerical model simulations to examine how, in general, RyR density affects the AP firing rate in SANC in different conditions. Intuitively, the higher density of the channels should always result in stronger Ca release and higher AP rates via the coupled-clock mechanism [4]. Model simulations, however, showed a counterintuitive result (Fig 6, red curve) that higher RyR densities decrease the AP rate (i.e. increase APCL). Our further examination of SR Ca dynamics (Fig 4D) revealed that an increase in RyR density (and the respective increase in junctional SR size accommodating the additional RyRs) shifts source-to-sink balance towards sink in Ca refilling of junctional SR from free SR. When sink increases, the same SR pumping capability (modest in the basal state) results in lower CaFSR levels. This leads, in turn, to slower CaJSR rise during diastolic depolarization and less LCR activity during diastolic depolarization (Fig 4C, scenario 2) and lower AP rate.

During βAR stimulation, SR pumping capability substantially increases via phospholamban phosphorylation [4] (parameter P_up_ in our model) with increased Ca influx via I_CaL_, naturally matching the increased Ca source to the increased sink (with larger RyR density). The SR Ca loading indeed increases in our model (Fig 4D) as shown in previous experimental studies [26]. This leads, in turn, to faster kinetics of junctional SR refiling with Ca (via diffusion from free SR), resulting in higher LCR activity during diastolic depolarization and pacemaker rate in the model (Fig 4, scenario 8 and Fig 6, blue curve) and previous experimental studies [26, 27]. Indeed, a CRU can generate a Ca spark when CaJSR reaches a critical level (300 μM in our model) that allows a phase transition of the RyR release channel system from a metastable state [28, 29]. This critical level is achieved the earliest during diastolic depolarization in the presence of βAR stimulation, explaining, in part, the highest LCR activity at the I_CaL_ threshold and the shortest APCL (Fig 4).

We also characterized the distribution of cluster sizes (RyRs per cluster), a parameter that had not been previously established. This parameter is important (along with the channel density) for Ca clock function, because activation and termination of a Ca spark critically depend on the number of release channels in the CRU [28–31]. In the literature, we found only preliminary results of dSTORM-based single RyR imaging in rat SANCs published by the Lederer’s group as abstracts [32, 33]. They found that RyRs organized into heterogeneous clusters ranging from 15 to 140 RyRs with short (often < 50 nm) edge-to-edge distances between adjacent clusters, suggesting a possibility of inter-cluster signaling. Our findings are consistent with the results of their study.

Hopkins statistic analysis reveals distinct changes in the internal spatial organization of RyRs within individual clusters for different cluster sizes under β-adrenergic stimulation (Fig 3). Superclusters (>76 RyRs) demonstrated a 6.2% increase in Hopkins values (p<0.01), though remaining below 0.4, suggesting a reorganization from grid-like uniformity toward a more heterogeneous internal architecture. This reorganization likely reflects the coalescence of adjacent RyR clusters into larger superclusters under βAR stimulation, as visualized in Fig 2. Coalescence of RyR clusters to form larger clusters has been also reported in ventricular myocytes after application of isoproterenol [24]. Under basal conditions (Fig 2B), individual clusters including a modest supercluster (99 RyRs) maintain relatively discrete boundaries with almost uniform internal RyR distribution. Following βAR stimulation (Fig 2D), massive superclusters (130, 188, and 414 RyRs) appear likely resulting from merger events, creating internal heterogeneity with dense RyR-rich cores separated by relatively sparse regions - a pattern consistent with the increased Hopkins values (∼0.33). This transition from uniform to heterogeneous organization within superclusters can be important to generate early and stronger synchronized release events (recruiting neighboring smaller CRU to fire, Fig 5) while avoiding metastability known for larger fully packed clusters (without voids) featuring stronger RyR interactions and generating noisy Ca leak [28]. Indeed, simulations of Ca sparks generated by RyR clusters of different geometries [34] showed that larger and denser clusters exhibit higher spark fidelity, i.e. the probability that a spontaneous RyR opening triggers a Ca spark. Therefore, less dense larger clusters such as found here (see also [24]) represent functional groupings of several neighboring clusters close enough that Ca diffusion can cooperatively recruit them into a single large spark while remaining less prone to metastability.

Our numerical simulations (Fig 4) isolated the functional impact of two distinct structural changes, density vs. clustering. The simulations revealed that RyR density and superclustering can be viewed as separate but synergistic regulatory mechanisms. Increasing RyR density alone prolonged APCL in the basal state (Fig 6). However, this high density became advantageous during βAR stimulation, where it worked with upregulated Ca² cycling and membrane ion currents to shorten the APCL (Fig 6).

Importantly, the “tail fraction” comprised of superclusters (Fig 2C) acted as a potent accelerator in all conditions. Introducing the supercluster distribution (from Cell 12) shortened the APCL under both basal (from 414.9 ms to 317.7 ms) and βAR-stimulated (from 284 ms to 269.1 ms) conditions by creating Ca² release “hotspots” that serve as nucleation sites (S1-S8 Videos). This two-tiered organization represents an efficient regulation mechanism, allowing SANCs to maintain a stable basal rate while retaining substantial reserve capacity by engaging higher-density networks and larger superclusters during βAR stimulation.

Our results on RyR clustering-superclustering provide a new dimension to the present perspective on RyRs as dynamic molecules that rearrange their location and function commensurate demand for Ca release under different conditions. RyR channels self organize into clusters whose size, packing, and spacing are highly plastic and adaptive to confer optimal CICR excitability. At the dyads, Junctophilin 2 and BIN1 arrange Cav1.2 and RyR2 into CRU super clusters [35, 36]. Acute β AR stimulation tends to enlarge CRUs and raise spark output [24, 37], whereas prolonged β AR or heart failure drives CaMKII/PKA dependent dispersion, fragmenting clusters and degrading CRU solidity [22, 38]. Luminal partners (CASQ2–triadin–junctin), accessories (FKBP), phosphorylation state, and the cytoskeleton fine tune this flexible architecture, giving the heart a dynamic range for beat to beat performance until disease shifts the balance [23, 25, 39, 40].

### Limitations and Future Studies

Our dSTORM analysis provides high-resolution structural snapshots but does not capture the dynamic behavior of RyR clusters. How specifically these clusters rearrange in time and space in response to βAR stimulation remains an important question that could be explored with live-cell super-resolution techniques. The findings are based on rabbit SANCs. While rabbit is a common model in cardiac pacemaker research, species differences in ion channel expression could differ substantially. Comparative studies in other species, including human SANCs, are needed to establish the generality of these findings. Any computational cell model is a simplification of biological reality. Our model incorporated the experimentally measured distributions of cluster sizes, but it does not capture all possible interactions of RyRs within CRUs and with other molecules in time and space. For example, while our CRU agent-based model of SANC can examine effects of RyR densities and cluster sizes, it does not simulate interactions of individual RyRs. Therefore, our speculation about importance of less organized clusters of larger sizes to avoid metastability will need further numerical validation using a more detailed model of CRUs (such as in [28, 34]) that include interactions of individual RyRs within the cluster. We only examined RyR network under the cell membrane. Future studies will address the question of how βAR stimulation affects RyRs in the cell interior and whether these changes are different from (or linked to) RyR distribution/density at the cell periphery. Our data analysis suggests that superclusters are likely formed by smaller cluster merger events. However, how specifically the clusters move, coalesce, expand, and join together in periphery regions remains to be investigated. Finally, further studies will clarify regional heterogeneity of supercluster distribution within cells that might relate to pacemaker site localization within SAN tissue.

### Clinical Relevance

Our finding that RyR superclusters are particularly important for achieving the highest possible heart rates during β-adrenergic stimulation is clinically relevant. Dysfunctional RyR gating and altered Ca handling are implicated in sinoatrial node dysfunction (SND) and arrhythmias, including sick sinus syndrome [41]. Cardiac aging is associated with a decline in pacemaker function and increased incidence of SND [42]. Our results suggest that disruptions or alterations in the nanoscale organization of RyR network (such as changes in cluster size distribution or density) could be an underappreciated mechanism contributing to SND. Here we show that alterations leading to less synchronized Ca release by a RyR network with fewer large clusters or excessive number of small, disorganized clusters could cause bradycardia that is known to culminate sinus arrest. Investigating how RyR cluster architecture changes with age using super-resolution microscopy could provide new insights into age-related SND.

Impaired β-adrenergic responsiveness is common in aging and heart failure. Our data suggests that structural alterations in the RyR network might contribute to this blunted chronotropic response. Therapeutic strategies aimed at preserving or restoring optimal RyR clustering could potentially improve heart rate regulation. For example, age-associated overexpression of BIN1 emerges alongside dysregulated endosomal recycling and disrupted trafficking of RyRs, whereas BIN1 knockdown restores the nanoscale distribution and clustering plasticity of cardiac RyRs [36].

Our results can be also important for sports medicine. Reaching the highest possible heart rate is often critical in setting new records of human abilities. On the other hand, athletes often have bradycardia at rest. It was shown that the training-induced bradycardia is not a consequence of changes in the activity of the autonomic nervous system but is caused by intrinsic electrophysiological changes in the sinoatrial node such as downregulation of HCN4 channels [43]. The present study can offer a new RyR network-related mechanism that explains the dual effect of exercise in athletes of reaching the highest possible heart rates at the expense of getting bradycardia at rest. Indeed, our results in Figs 4 and 6 demonstrate that increase in RyR density could lead to lower pacemaker rates at rest yet yielding higher rates during exercise.

Understanding the importance of RyR clustering might open new therapeutic avenues. While directly manipulating nanoscale protein organization is challenging, strategies aimed at modulating factors that influence RyR clustering or stabilizing functional cluster sizes could potentially be explored for treating SND or improving heart rate control. Further research correlating RyR nanoscale structure with functional readouts in relevant clinical models (e.g., aging animals, models of heart failure) is needed to fully explore these implications.

### Conclusions

Our study utilized super-resolution dSTORM microscopy to reveal, for the first time, the detailed nanoscale organization of ryanodine receptors beneath the membrane of rabbit SANCs. We discovered significant heterogeneity in RyR density and demonstrated that RyRs form clusters with a right-skewed size distribution, characterized by the prevalence of numerous small clusters alongside a smaller subset of large, functionally significant superclusters. Our computational modeling indicated that this specific architecture, particularly the proportion of large clusters, critically determines the efficiency of local Ca release and thereby modulates SANC automaticity. Larger clusters facilitate faster pacemaking under β-adrenergic stimulation by firing earlier during diastole and recruiting smaller clusters to fire thereby timely synchronizing Ca release among cluster populations of different sizes. Overall, our new mechanism of β-adrenergic stimulation proposes that a substantial population of smaller RyR clusters do not fire during diastolic depolarization but stay in “reserve” to be recruited during stimulation as an additional diastolic Ca signal source that accelerates diastolic depolarization and AP firing rate. Our findings underscore the crucial role of RyR nanoscale organization, in synergy with SR Ca pumping and cell membrane ion channels (I_CaL_, I_Kr_, and I_f_), as a powerful mechanism regulating cardiac pacemaker function and adaptability.

## Materials and methods

### Enzymatic isolation of individual SANC

SANCs were isolated from male rabbits in accordance with NIH guidelines for the care and use of animals, protocol # 457-LCS-2024, as previously described [44]. New Zealand white rabbits (Charles River Laboratories, USA) weighing 2.8–3.2 kg were anesthetized with sodium pentobarbital (50–90 mg/kg). The hearts were removed quickly and placed in solution containing (in mM): 130 NaCl, 24 NaHCO_3_, 1.2 NaH_2_PO_4_, 1.0 MgCl_2_, 1.8 CaCl_2_, 4.0 KCl, and 5.6 glucose equilibrated with 95% O_2_/5% CO_2_ (pH 7.4 at 35 °C). The sinoatrial node region was cut into small strips (∼1.0 mm wide) perpendicular to the crista terminalis and excised. The final sinoatrial node preparation, which consisted of sinoatrial node strips attached to the small portion of crista terminalis, was washed twice in nominally Ca-free solution containing (in mM) 140 NaCl, 5.4 KCl, 0.5 MgCl2, 0.33 NaH_2_PO_4_, 5 HEPES, and 5.5 glucose (pH = 6.9) and incubated on a shaker at 35 °C for 30 min in the same solution with the addition of elastase type IV (0.6 mg/mL; Sigma, Chemical Co.), collagenase type 2 (0.8 mg/mL; Worthington, NJ, USA), Protease XIV (0.12 mg/mL; Sigma, Chemical Co.), and 0.1% bovine serum albumin (Sigma, Chemical Co.). The sinoatrial preparation was next placed in modified “Kraftbruhe” solution, containing (in mM) 70 potassium glutamate, 30 KCl, 10 KH_2_PO_4_, 1 MgCl_2_, 20 taurine, 10 glucose, 0.3 EGTA, and 10 HEPES (titrated to pH 7.4 with KOH), and kept at 4 °C for 1 h in the Kraftbruhe solution containing 50 mg/mL polyvinylpyrrolidone (Sigma, Chemical Co.). Finally, cells were dispersed from the sinoatrial node preparation by gentle pipetting in the “Kraftbruhe” solution and stored at 4 °C.

### Staining and mounting

Isolated rabbit SANCs were plated on #1 glass, 14-mm glass-bottom MatTek dishes pre-coated with laminin (∼40 µg/mL) in phosphate-buffered saline (PBS) with 1% penicillin/streptomycin. Cells were rinsed twice with PBS, fixed in 2–4% paraformaldehyde for 10 min at room temperature, rinsed, quenched in 100 µM glycine for 10 min, rinsed again, permeabilized with 1% Triton X-100 in PBS for 15 min, and rinsed twice with 0.1% Triton/PBS. Samples were then blocked in PBS containing 2% IgG-free BSA, 5% normal goat serum, 0.02% sodium azide, and 0.1% Triton X-100 for 60 min to 4 h. Our primary antibodies were RyR2 monoclonal antibodies (clone C3-33, Thermo Fisher Scientific cat# MA3-916). The antibodies diluted (1:100) in blocking buffer were applied for 60 min at room temperature or overnight at 4 °C, followed by four 5–10 min washes in 0.1% Triton/PBS. Then cross-adsorbed secondary antibody F(ab’)2-Goat anti-mouse IgG (H+L), Alexa Fluor™ 647 (Thermo Fisher Scientific cat# A-21237) diluted 1:200 were applied to cells and incubated for 60 min at room temperature or overnight at 4 °C in the dark, followed by four 5–10 min washes in 0.1% Triton/PBS in the dark and two final PBS rinses. Then the Alexa Fluor 647-labeled SANCs were submersed in an imaging buffer containing 20% VectaShield (H-1000, Vector Laboratories) diluted in Tris-glycerol (5% v/v TRIS 1 M pH 8 in glycerol, Sigma-Aldrich. For experiments involving acute βAR stimulation, before fixation SANC were equilibrated 15–20 min at 37 °C in physiological solution containing 1.8 mM Ca², then treated with 0.3 µM isoproterenol for 5 min at 37 °C; control cells received vehicle only.

### Single-molecule detection

The prepared and mounted SANC samples were imaged using ZEISS Elyra 7 Super resolution microscope, equipped with dSTORM single molecule localization microscopy module (SMLM). Fluorophore emission light was collected via an alpha Plan-Apochromat 100× objective lens and Immersol 518 F fluorescence-free immersion oil (NE = 1.518 at 23°C). The laser beam was adjusted to TIRF incidence to selectively excite fluorophores on the cell surface within approximately 100 nm of the coverslip. Blinks were collected across a 512 × 512 μm field of view on the cell surface. All original dSTORM data in the form of Raw SMLM Molecule Tables are available online at Harvard Dataverse https://doi.org/10.7910/DVN/T0REJ3.

Single-molecule detections were performed using ZEN Black 3.0 software throughout each acquisition. The peak mask size was set to 9 pixels, defining the window around each candidate molecule that was isolated for fitting. A peak intensity-to-noise ratio threshold of 6 was applied, so that only peaks at least sixfold above the local background were accepted as genuine detections. Each isolated peak was then fit with the software’s point-spread-function model to obtain subpixel coordinates, and positions were stored as floating-point values (rounded to the nearest nanometer for downstream analysis). Subsequent quality filters and analyses were applied as described in the following sections.

### Image analysis

Our Super-Resolution Single-Molecule Localization Microscopy (SMLM) processing pipeline, depicted in the flowchart in Fig 7 used ZEN Black 3.0 software application supplemented with our Python and MATLAB code (available at GitHub : https://github.com/valventura/dSTORM). The pipeline performs precise filtering, cell region delineation, clustering, and statistical analysis of RyR localizations in SANCs. Our analysis is divided into two major stages: (1) pre-processing and filtering of the *Raw SMLM Molecule Table* generated by the ZEN black software and (2) clustering and statistical analysis of this final particle table.

**Fig 7.**
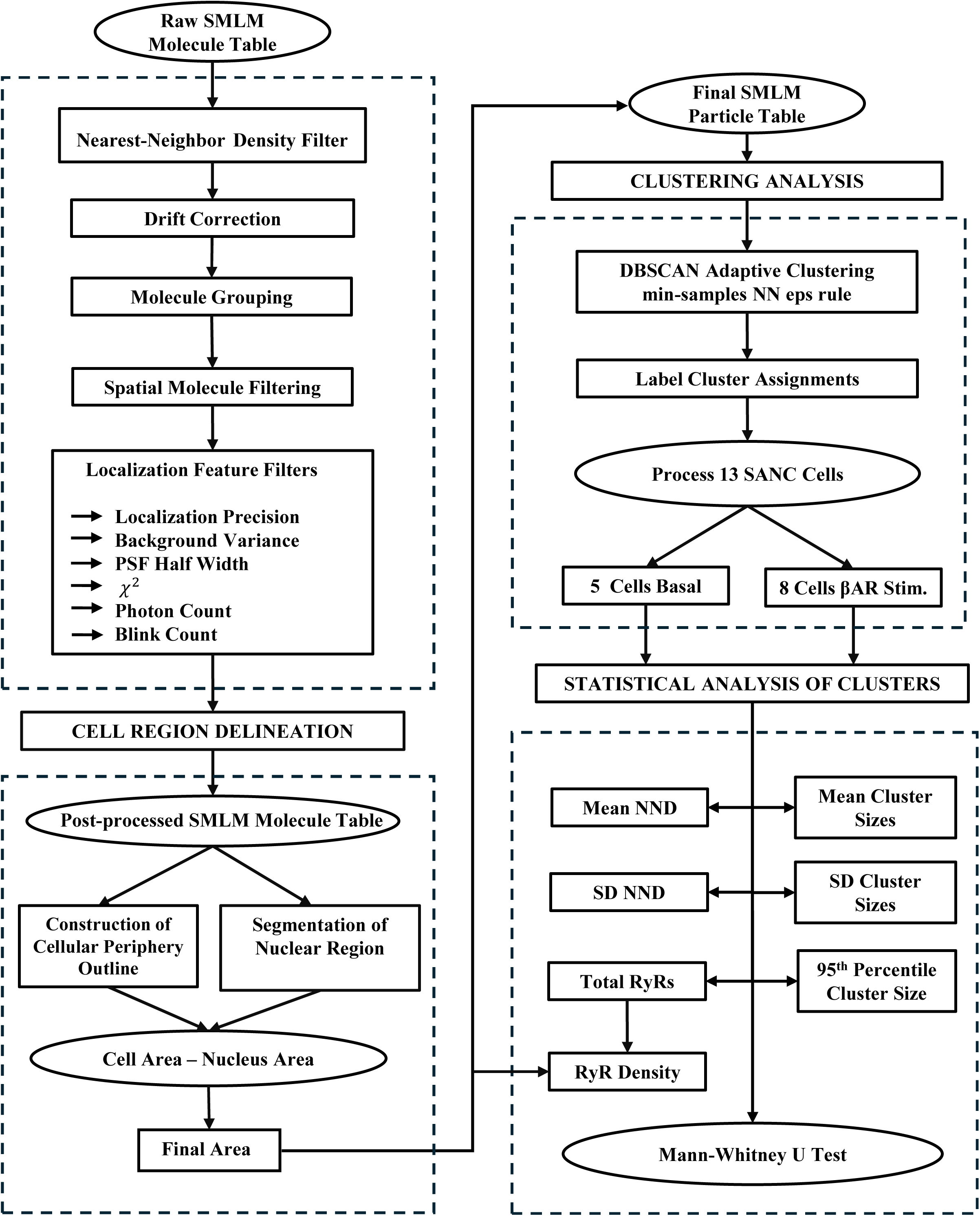
Flowchart of the dSTORM-based RyR cluster analysis algorithm. Presented is our Single-Molecule Localization Microscopy (SMLM) processing pipeline for precise filtering, delineation, clustering, and statistics of RyR localizations in SANCs. Raw detections are drift-corrected, grouped, and spatially filtered, then gated by localization precision (1–30 nm), photon count (1,000–6,000), PSF half-width (6.42–250 nm), χ² (0–2), and background (≤ mean + 1.5·SD), with a ≥6 blinks-per-site rule. Cell region is defined by a periphery mask with nuclear-void removal (if applicable) to compute the final area. The post-processed particles are clustered by DBSCAN using an adaptive nearest-neighbor ε; from the labels we derive nearest neighbor distances (NNDs), cluster-size metrics (mean, SD, 95th percentile), and RyR density, and compare groups by two-sided Mann–Whitney U tests.

#### SMLM localization filtering and delineation

Raw SMLM detections were subjected to a multi-stage filtering process to remove low-quality localizations. An example of filtering within cell area and related histograms are shown in Figs 8 and 9, respectively.

**Fig 8.**
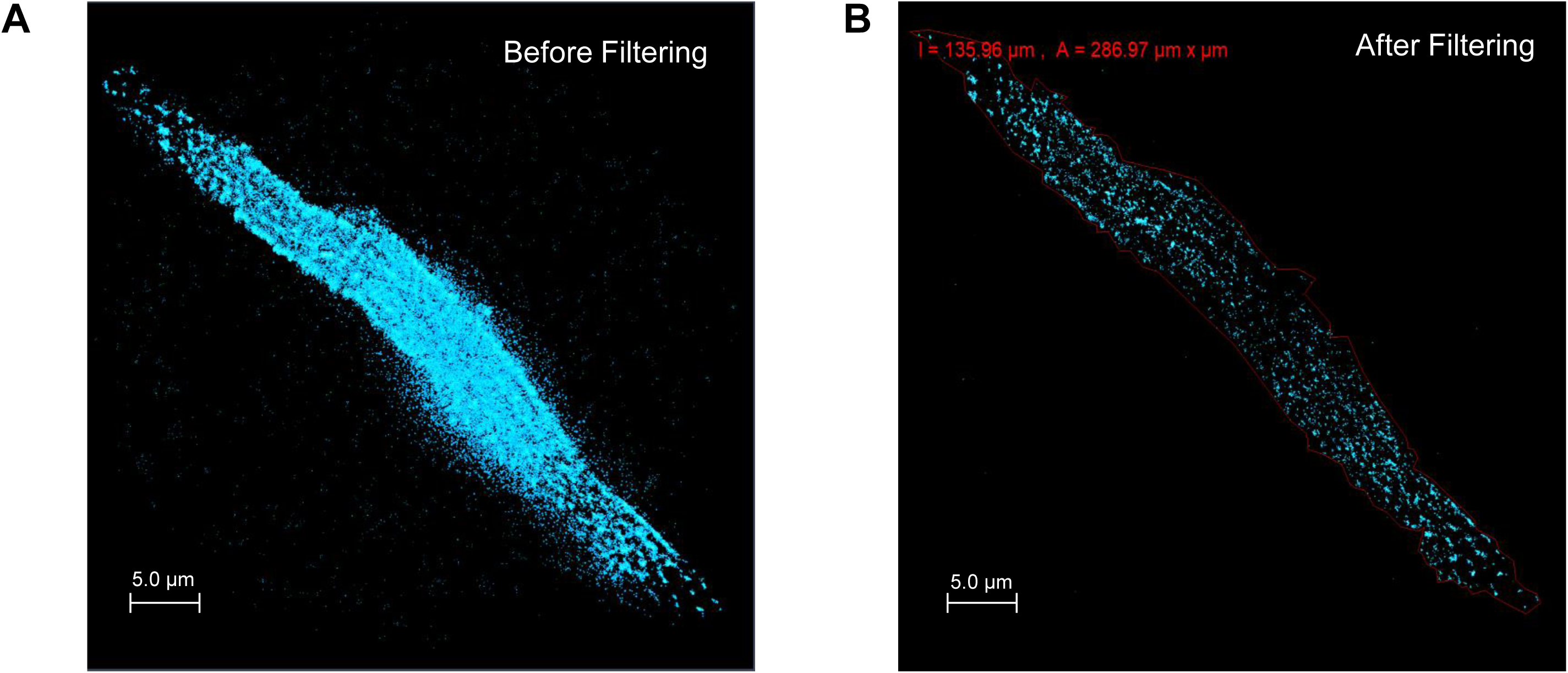
Filtering of RyR dSTORM localizations in a SANC. Shown is a representative cell before and after the filtering pipeline. **A.** Unfiltered localizations (cyan). **B.** Retained localizations (cyan) after applying the periphery mask (red polygon) together with automated windows on localization quality and spatial context. Colors/symbols: cyan dots are RyR events; red polygon is the cell periphery; the red header reports the periphery perimeter (l = 135.96 µm) and area (A = 286.97 µm^2^). Filters applied: precision 1–30 nm, photons 1,000–6,000, PSF half-width 6.42–250 nm, χ² 0–2, background < mean + 1.5 SD; and a blinks-per-site rule (≥6 molecular blinks within 50 nm). Together, panels A–B demonstrate removal of extra-cellular and low-quality detections while preserving the in-cell high-quality RyR localization detections.

**Fig 9.**
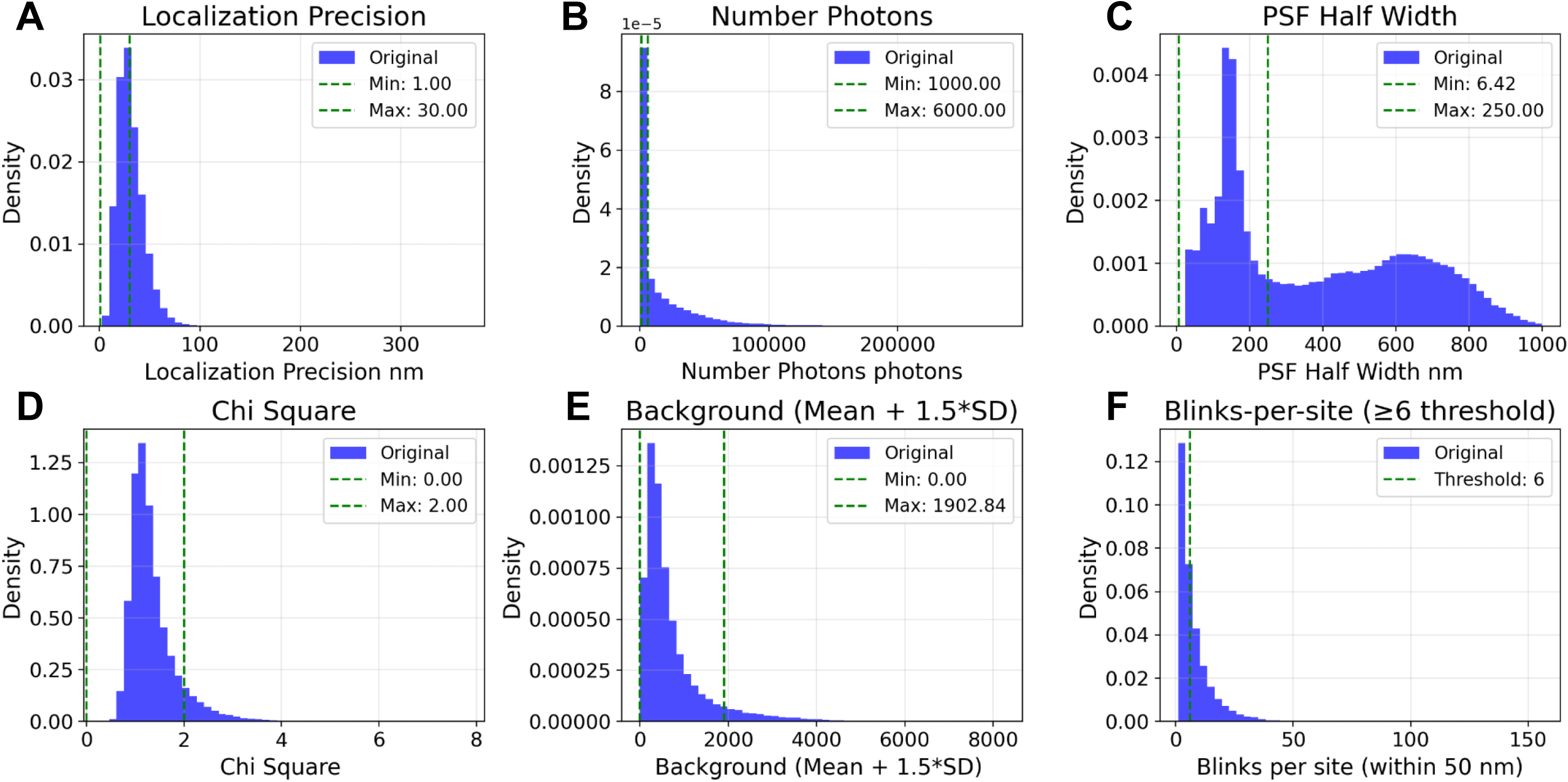
Distributions of dSTORM localization parameters and their filtered values. Density histograms (blue) of the raw localization parameters (calculated for cell 6); green dashed lines mark the acceptance limits used in the filtering pipeline. **A–E.** Filter gates are applied to all detections: localization precision 1–30 nm, number of photons 1,000–6,000, PSF half width 6.42–250 nm, χ² 0–2, and background capped by a dynamic limit of mean + 1.5·SD (0–1902.84 in this dataset). These windows are indicated by the dashed vertical lines in each panel. **F.** Distribution of blinks-per-site computed in a 50 nm neighborhood; the dashed line marks the ≥6 blinks threshold used to define stable RyR sites. Together, the figure summarizes the criteria that remove low-precision, low-photon, poorly fit, or high-background events while preserving high-quality in-cell detections; across this cell, 111,307 original localizations were reduced to 13,894 after all gates including the ≥6 blinks rule (12.5% retained).

##### 1. Nearest-Neighbor Density Filter

Each localization was evaluated within a circular search area, and if the number of neighboring peaks was below the position threshold, that localization is considered an outlier and removed from the SMLM image and molecule list. For each localization, the number of neighboring molecules within a radius of 100 nm was counted; localizations with fewer than two neighbors were excluded. Single or couples of isolated RyRs likely represent either non-functional receptors, RyRs not yet incorporated into functional release sites, or artifacts from antibody labeling and imaging.

##### 2. Drift Correction

Sample drift was corrected with the Model based method in ZEN Black 3.0. In this approach, the software estimates a linear drift directly from the sample structures. For each segment of the acquisition, a correction vector is computed from the average alignment of all structures. We set Correction Type = Model based and enabled segmentation with Segments = Automatic while limiting the max to 4. In automatic mode, the algorithm chose the number of segments but never exceeded the user defined maximum. The drift was first corrected within each segment and then residual offsets between segments were corrected.

##### 3. SMLM Grouping

The localization data were grouped in ZEN Black 3.0 using the *SMLM-Grouping* tool to merge repeated detections of the same molecule over consecutive frames. Grouping parameters were chosen Max On Time = 5 frames, Off Gap = 10 frames, and Capture Radius = 1.0 pixels.

The algorithm identifies peaks that fall within the defined capture radius across successive frames and assigns them to a single molecule if the temporal constraints are met.

- *Max On Time* limits the maximum number of consecutive frames in which a fluorophore can appear while still being treated as one molecule. Peaks persisting longer than this are discarded to prevent false merging of overlapping emitters.
- *Off Gap* specifies how many frames the fluorophore may temporarily disappear while still being linked to the same molecule. If this limit is exceeded, the peaks are separated into distinct molecules.
- *Capture Radius* defines the spatial tolerance [set to 1.0 pixel (x, y)] within which peaks are considered part of the same molecule based on the rendered localization precision.

After applying these parameters, the Group function was executed to merge localizations meeting the criteria. This procedure reduced redundant detections from fluorophore blinking, yielding a refined SMLM molecule table with fewer but more accurately positioned molecular localizations.

##### 4. Localization Feature Filters

We applied a series of acceptance windows based on the localization fit parameters, as summarized in S2 Table. These gates included:

- Localization Precision: 1.0–30.0 nm
- Photon Count: 1,000–6,000 photons
- PSF Half Width: 6.42–250.0 nm
- Chi Square(χ^2^): 0.0–2.0
- Background Variance: A dynamic threshold was applied, retaining localizations with background variance < (*mean* = 1.5 x *SD)* of the cell’s total background distribution.

##### 5. Cell Region Delineation

We analyzed data only within cell perimeter, i.e. falling outside the periphery polygon (Fig 8). In some images, depending on proximity to cell surface, nuclei clearly influenced RyR detection, and their respective regions were also excluded from analysis (“Cell Area - Nucleus Area”).

##### 6. Site Stability Filter

To ensure that retained localizations represent stable RyR sites, we combined a spatial neighborhood check with a blink-based stability criterion. First, using a k-d tree, we counted for each localization the number of neighboring localizations within a 50 nm radius. We then computed the number of detected blinks (blinks_per_site) for each site as the total count of temporally linked events assigned to that position. Sites exhibiting fewer than six blinks were excluded from further analysis, an empirically determined threshold indicative of stable fluorophore behavior [45]. A similar filtering algorithm (≥5 neighbors within 60 nm) was used previously in [46]. This two-step filter removed low-confidence, transient detections and retained reproducible RyR localizations (Fig 9).

The set of localizations passing all features, stability, and spatial gates constituted the *Post-processed SMLM Molecule Table*, which was used for all subsequent clustering.

#### Clustering analysis

The RyR localizations from the *Post-processed SMLM Molecule Table* were clustered using the Density-Based Spatial Clustering of Applications with Noise (DBSCAN) algorithm to identify

RyR clusters. The two key parameters for DBSCAN, min_samples and ε, were determined based on the biophysical characteristics of RyR clusters and the properties of SMLM data.

We defined cluster cores using a min_samples of 3. Given that functional RyR clusters (CRUs) can be small, this value was selected as the minimal robust definition of a dense region (a point and two neighbors). This low threshold provided the necessary sensitivity to detect the smallest biologically relevant RyR aggregates, which may consist of only a few RyR molecules. A higher threshold (e.g., 5 or 10) would risk misclassifying these small but significant nascent clusters as noise, thereby skewing the resulting size and density distributions.

Our calculation of *clustering radius (*ε) in our DBSCAN algorithm is as follows:

1. For a given cell with *N* localizations, the distance *d_i_* to the 3rd nearest neighbor (i.e., corresponding to min_samples = 3) was computed for each point. *i*.
2. From the resulting set of *N* distances *{d_i_*} we identified all positive distances *d_i_ >0* and computed their 5^th^ percentile *P*.
3. The clustering radius *ε* is defined as

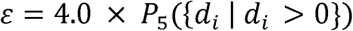

*ε* functions as a dimensionless interaction envelope: large enough to preserve intra-cluster cohesion across density gradients, yet small enough to maintain inter-cluster separation. An example result of DBSCAN clustering is shown in Fig 10.

**Fig 10.**
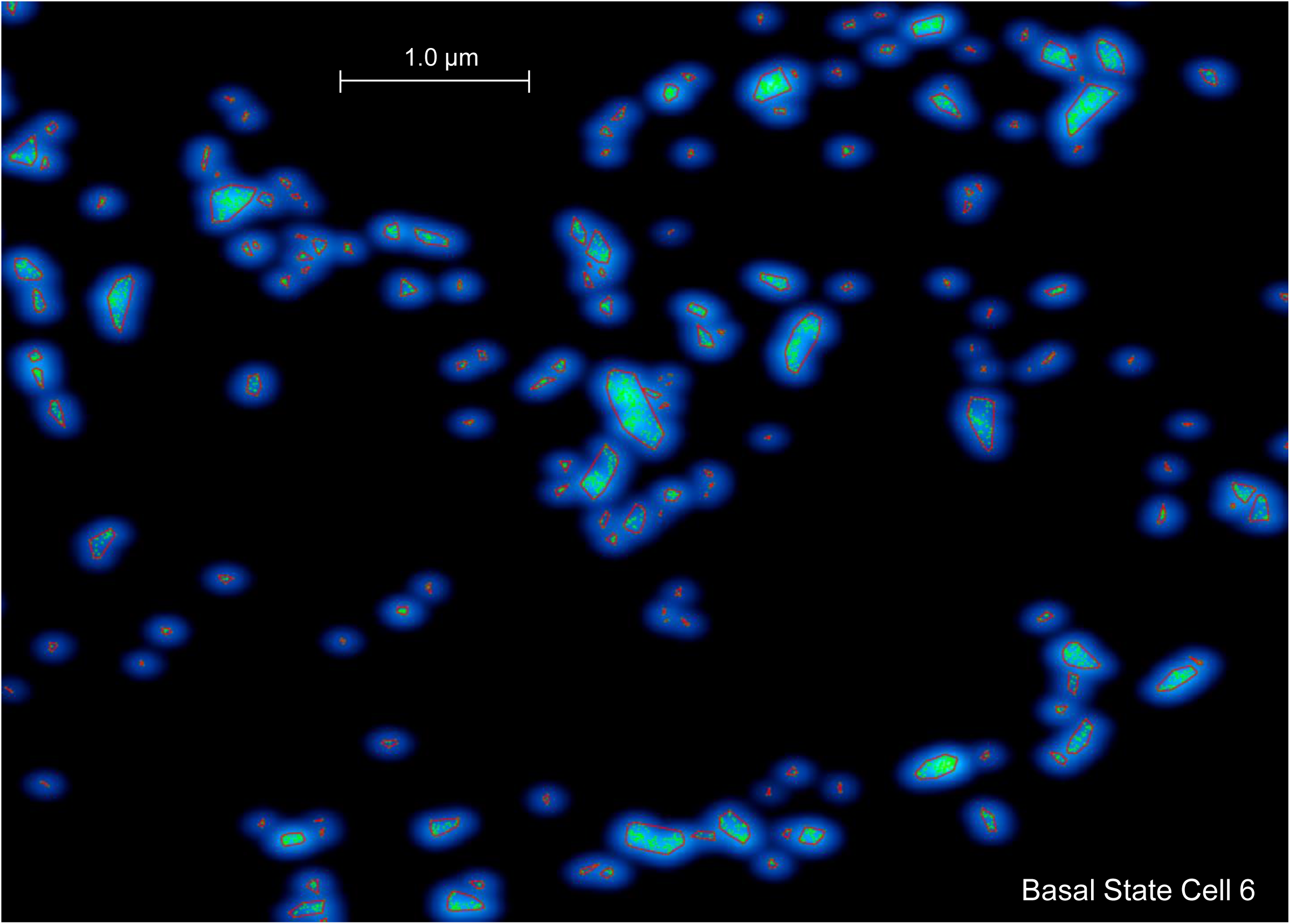
DBSCAN clustering (red outlines) of RyR dSTORM localizations in a basal SANC. A representative example (Cell 6) showing adaptive DBSCAN labels on retained localizations. A smoothed “blue-glow” density map forms the backdrop, lime dots mark RyR events, and red polygons trace convex-hull boundaries of individual clusters. Colors/overlays derive from the plotting routine (blue gradient heatmap + lime points; red hulls). DBSCAN used the min-samples-th nearest-neighbor ε rule (ε = 4.0 × 5th percentile of positive min_samples-th NN distances) with min_samples = 3; points labeled as noise are omitted from the display.

#### Cluster geometry and spacing analysis

Following DBSCAN, we quantified the geometry and spatial relationships of all resulting clusters.

1. Cluster Size and Density: Cluster size was defined as the total number of localizations (RyR) assigned to that cluster label. We computed the mean, standard deviation (SD), and 95th percentile of cluster sizes on a per-cell basis. RyR Density (RyR/μm^2^) was calculated as the total number of clustered localizations divided by the final area.
2. To determine cluster spacing (nearest neighbor distance), we first computed the 2D convex hull for every individual cluster. A global k-d tree was then constructed from all vertices of all hulls in the cell. For each cluster we found the shortest Euclidean distance from any of its hull vertices to a vertex belonging to any other cluster. This shortest inter-hull distance was recorded, and the per-cell Mean and SD of nearest neighbor distance were calculated from this set of inter-cluster distances. Our measured nearest edge-to-edge distances have physiological relevance as RyR clusters interact via CICR mechanism activated when released Ca from one cluster reaches RyRs at the border of a neighboring cluster.
3. Statistical Analysis: Comparisons between basal and βAR-stimulated groups for all derived metrics were performed using a two-sided Mann-Whitney U test.

#### Hopkins statistic calculation

To quantify the internal spatial organization of RyRs within individual clusters, we calculated the Hopkins statistic with 100 iterations per cluster to ensure statistical stability. The Hopkins statistic (H) tests the spatial randomness hypothesis by comparing distances from random points to their nearest data points with distances between data points themselves:

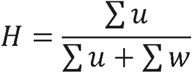

where u represents the distance from each randomly selected RyR to its nearest neighbor within the cluster (excluding itself), and w represents the distance from each randomly generated point within the cluster boundary to its nearest RyR. Both sums are taken over n sampled points. For each cluster, we sampled n points where n = min(max(10, N/10), 30) and N is the total number of RyRs in the cluster. This ensures a minimum of 10 samples and a maximum of 30 samples to balance computational efficiency with statistical power. Clusters containing fewer than 10 RyRs were excluded from Hopkins analysis due to insufficient points for reliable calculation.

The Hopkins statistic ranges from 0 to 1 with the following interpretations:

H ≈ 0.5 indicates spatially random distribution (Poisson process)

H → 1 indicates clustered/aggregated distribution (points form clumps within the cluster boundary)

H → 0 indicates regular/over-dispersed/inhibitory pattern (points are more evenly spaced than random, i.e., “anti-clustered”)

Statistical comparisons between basal and βAR conditions were performed using Mann-Whitney U tests for each size category.

### Numerical model simulations

Numerical simulations were performed using a new SANC model featuring a functional network of individual RyRs. We developed this model based on our previous agent-based model in which interacting agents were CRUs [14]. Due to voxel size limitations, the CRU sizes in that old model were constructed as multiples of 16 RyRs. In the new model used in the present study, we overcame this limitation. In the new model, the CRUs sizes (and their reciprocal junctional SR sizes and Ca releases) were constructed with single RyR precision to closely reproduce realistic RyR cluster network obtained by our dSTORM imaging of individual RyR molecules. The new model code is provided at GitHub: https://github.com/victoramaltsev/RyR-network-SANC-model. The CRUs were randomly rotated and (uniformly) randomly distributed under cell membrane. To minimize effect of randomness on the simulation results, we simulated data for five different random realizations for each simulation scenario of 8 (total 40 simulation runs of 16 s duration) with average data presented in Fig 4B. In each simulation run, APCL data were analyzed for the last 10 s, when the system achieved a steady firing pattern. βAR stimulation was simulated in the model with respective change in the model parameters describing SR Ca pumping rate (P_up_) and ion currents as previously reported [14].

## Supporting information

**S1 Table. Per-cell RyR density, cluster size, and spacing metrics from DBSCAN clustering.** (MS Word)

**S2 Table. Acceptance windows for dSTORM localization quality and site stability. (MS Word)**

**S1-S8 Videos** (MP4). 2D Ca dynamics under cell membrane during the last second of each 16 s simulation, with video number corresponding to simulation scenario number from 1 to 8:

1. Basal, Normal Density, Normal Clustering
2. Basal, High Density, Normal Clustering
3. Basal, Normal Density, Superclustering
4. Basal, High Density, Superclustering
5. βAR, Normal Density, Normal Clustering
6. βAR, High Density, Normal Clustering
7. βAR, Normal Density, Superclustering
8. βAR, High Density, Superclustering

“Basal” designates cell model parameters reflecting pacemaker cell operation in the basal state. “βAR” designates enhanced model parameters P_up_ (SR Ca pump), I_CaL_, I_Kr_ and I_f_ reflecting βAR stimulation. “Normal Density” was set to 67.65 RyR/μm^2^, the average density in the basal state and “High Density” was set to 119.07 RyR/μm^2^, the average density during βAR stimulation (S1 Table). “Normal Clustering” was set to RyR cluster size distribution in Cell 6 (basal state) and “Superclustering” was set to RyR cluster size distribution in Cell 12 (βAR stimulation). [Ca] was coded in videos by red shades from black (0.15 μM) to pure red (>10 μM). CRU functional states were coded by colors as follows: refractory CRUs are in blue shades; CRUs ready to fire are in green; and CRUs releasing Ca are in grey shades. Both blue shades and white shades reflect JSR Ca changes, with a saturation level set at 300 µM (Ca spark activation threshold in our model). Simulation time and membrane potential are shown at the bottom left corner of each video.

## Acknowledgments

The authors thank Mr. Bruce Ziman for skillful isolation of sinoatrial node cells.

## Data availability statement

Our data analysis used ZEN Black 3.0 software (Zeiss company) supplemented by our Python and MATLAB. The supplemental analysis code is available at GitHub: https://github.com/valventura/dSTORM. The code for our new numerical model is available at GitHub: https://github.com/victoramaltsev/RyR-network-SANC-model. Original data used in our analysis are available online at Harvard Dataverse https://doi.org/10.7910/DVN/T0REJ3.

## Author contributions

**Conceptualization:** Victor A Maltsev, Michael D Stern, Edward G Lakatta

**Formal analysis:** Valeria Ventura Subirachs, Alexander V. Maltsev, Syevda Tagirova, Victor A Maltsev

**Funding acquisition:** Michael D Stern, Edward G Lakatta

**Investigation:** Syevda Tagirova, Valeria Ventura Subirachs, Dongmei Yang

**Methodology:** Valeria Ventura Subirachs, Syevda Tagirova, Dongmei Yang, Victor A Maltsev

**Software:** Valeria Ventura Subirachs, Alexander V. Maltsev, Victor A Maltsev

**Supervision:** Michael D Stern, Victor A Maltsev

**Visualization:** Valeria Ventura Subirachs, Alexander V. Maltsev

**Writing – original draft:** Valeria Ventura Subirachs, Victor A Maltsev

**Writing – review & editing:** Valeria Ventura Subirachs, Victor A Maltsev, Michael D Stern, Edward G Lakatta

## List of acronyms

AP: action potentials
APCL: AP cycle length
BIN1: Bridging Integrator 1
CaMKII: Calcium/Calmodulin-Dependent Protein Kinase II
CICR: Ca-induced-Ca-release
dSTORM: direct Stochastic Optical Reconstruction Microscopy
EGTA: Ethylene glycol bis(β-aminoethyl ether)-N,N,N’,N’-tetraacetic acid
HEPES: 4-(2-hydroxyethyl)-1-piperazineethanesulfonic acid
I_CaL_: L-type Ca current I_f_ funny current
I_Kr_: rapid delayed rectifier potassium current
LCR: local Ca releases
NCX: Na/Ca exchanger
NND: nearest neighbor distance
PBS: phosphate-buffered saline
PKA: cAMP-dependent protein kinase A
P_up_: Maximum SR Ca pumping rate in our model
RyR: ryanodine receptor
SANC: sinoatrial node cell
SERCA: Sarco/Endoplasmic Reticulum Calcium-transporting ATPase
SND: Sinoatrial node dysfunction SR sarcoplasmic reticulum
βAR: β-adrenergic receptor

