## Supplementary material for "A New Fight-or-Flight Pacemaker Mechanism via Ryanodine Receptor Abundance and Superclustering": S1 Table

|  | **Cell** | **RyR density, RyR·µm⁻²** | **Mean cluster size (RyR)** | **SD cluster size (RyR)** | **95th percentile cluster size (RyR)** | **Mean NND (nm)** | **SD NND (nm)** |
| --- | --- | --- | --- | --- | --- | --- | --- |
| **Basal state** | 1 | 96.12 | 26.98 | 65.24 | 90.00 | 159.90 | 145.03 |
|  | 2 | 16.44 | 10.52 | 11.03 | 28.65 | 303.82 | 272.71 |
|  | 3 | 47.23 | 14.83 | 20.37 | 42.55 | 185.82 | 174.83 |
|  | 4 | 71.08 | 16.27 | 30.14 | 49.00 | 149.27 | 172.41 |
|  | 5 | 123.82 | 20.49 | 30.93 | 66.40 | 119.57 | 121.27 |
|  | 6 | 51.24 | 15.35 | 16.99 | 47.95 | 162.53 | 167.68 |
|  | **AVG** | **67.65** | **17.40** | **29.11** | **54.09** | **180.15** | **175.65** |
| **βAR stimulation** | 7 | 161.12 | 23.94 | 33.92 | 88.00 | 103.30 | 100.49 |
|  | 8 | 122.14 | 22.39 | 35.91 | 82.70 | 104.58 | 120.25 |
|  | 9 | 92.78 | 18.69 | 23.93 | 59.95 | 150.04 | 136.79 |
|  | 10 | 90.96 | 20.82 | 27.79 | 71.10 | 136.24 | 132.26 |
|  | 11 | 156.61 | 28.09 | 57.37 | 120.65 | 100.16 | 117.67 |
|  | 12 | 120.82 | 22.23 | 34.77 | 86.00 | 109.43 | 124.79 |
|  | 13 | 125.70 | 22.47 | 31.12 | 80.00 | 109.15 | 128.82 |
|  | 14 | 82.71 | 22.67 | 39.18 | 71.00 | 131.22 | 167.14 |
|  | **AVG** | **119.11** | **22.66** | **35.50** | **82.43** | **118.02** | **128.53** |
|  | **% change** | **+76.10%** | **+30.23%** | **+22.00%** | **+52.40%** | **-34.49%** | **-36.66%** |

**S1 Table. Per-cell RyR density, cluster size, and spacing metrics from DBSCAN clustering.** Listed are values for cells in basal state (1–6) and βAR-stimulated cells (7–14): RyR density, mean cluster size with SD, 95th-percentile cluster size, and mean NND with SD. Group averages (AVG) are also shown; the bottom row reports the percent change (βAR vs. basal). Abbreviations: βAR, β-adrenergic receptor; NND, nearest-neighbor distance; SD, standard deviation; RyR, ryanodine receptor; DBSCAN, density-based spatial clustering of applications with noise.
