## Supplementary material for "A New Fight-or-Flight Pacemaker Mechanism via Ryanodine Receptor Abundance and Superclustering": S2 Table

| **Localization Feature** | **Lower bound** | **Upper bound** | **Description** |
| --- | --- | --- | --- |
| **Localization Precision** | **1.0** | **30.0** | **Calculated localization precision of a peak/object in nm in x and y.** |
| **Background Variance** | **0.0** | **Mean + 1.5 SD** | **Variance of the number of background photons.** |
| **PSF Half Width** | **6.42** | **250.0** | **Width of the point spread function (PSF) from the Gauss fit in x and y.** |
| **Chi Square (χ²)** | **0.0** | **2.0** | **χ² value of the Gauss fit for the localization precision.** |
| **Photon Count** | **1000.0** | **6000.0** | **Total number of photons detected from a peak/object.** |
| **Blink Count** | **6.0** | **∞** | **Spatial blinking density: number of co-localized detection events (within a 50 nm neighborhood).** |

**S2 Table. Acceptance windows for dSTORM localization quality and site stability.** Listed are the bounds used in the filtering pipeline and shown as dashed limits in Fig 9A–F. Together these limits remove low-precision, low-photon, poorly fit, or high-background noise events while preserving high-quality in-cell-perimeter RyR detections.
